# Different activation signatures in the primary sensorimotor and higher-level regions for haptic three-dimensional curved surface exploration

**DOI:** 10.1101/2020.08.04.235275

**Authors:** Jiajia Yang, Peter J. Molfese, Yinghua Yu, Daniel A. Handwerker, Gang Chen, Paul A. Taylor, Yoshimichi Ejima, Jinglong Wu, Peter A. Bandettini

## Abstract

Haptic object perception begins with continuous exploratory contacts, and the human brain needs to accumulate sensory information continuously over time. However, it is still unclear how the primary sensorimotor cortex (PSC) interacts with these higher-level regions during haptic exploration across time. This functional magnetic resonance imaging (fMRI) study investigates time-dependent haptic object processing by examining brain activity during haptic 3D curve and roughness estimation. For this experiment, we designed sixteen haptic stimuli (4 kinds of curve × 4 kinds of roughness) for the haptic curve and roughness estimation tasks. Twenty participants were asked to move their right index and middle fingers along with the surface twice and to estimate one of the two features--roughness or curvature--dependent on the task instruction. We found that the brain activity in several higher-level regions (e.g., bilateral posterior parietal cortex) linearly increased with curvature through the haptic exploration phase. Surprisingly, we found that the contralateral PSC was parametrically modulated by the number of curves only during the late exploration phase, but not during the early exploration phase. In contrast, we found no similar parametric modulation activity patterns for haptic roughness estimation in either the contralateral PSC or in the higher-level regions. Together, our findings suggest that haptic 3D object perception is processed across the cortical hierarchy, while the contralateral PSC interacts with other higher-level regions across time in a manner that is dependent upon object features.

**Highlights:**

- We observed the brain activity of haptic object perception using parametric stimuli.
- Haptic curve estimation showed parametric modulation across the cortical hierarchy.
- Curve parametric modulation in the sensorimotor cortex showed time dependency.
- Roughness parametric modulation showed very little dependency in any regions of the brain.
- These findings reflect the nature of time-dependent haptic object processing in the brain.

## 1. Introduction

In the somatosensory system, haptic perception originates through continuous exploratory contact with objects, and the human brain has to accumulate sensory information continuously across time to understand the object by touch (Klatzky and Lederman, 2011). During haptic exploration, both cutaneous and proprioceptive information are known to first arrive at the contralateral primary sensorimotor cortex (PSC) in the cerebral cortex (Pleger and Villringer, 2013; Sathian, 2016). Both non-human primate (NHP) (Arce-McShane et al., 2016; Umeda et al., 2019) and human neuroimaging (Huber et al., 2017) studies have demonstrated that the primary somatosensory cortex (S1) and the primary motor cortex (M1) are interacting with each other to shape the haptic information at the early stage. After such initial sensorimotor processing in the PSC, the integrated representation of the object local (e.g., surface roughness) and global (e.g., three-dimensional (3D) shape) features will be sent to other higher-level regions for further processing (Ackerley and Kavounoudias, 2015). However, the questions of whether and how haptic information are updated across time through the cortical hierarchy remain poorly understood.

S1 is known to comprise four cytoarchitectonic areas (areas 3a, 3b, 1, and 2), which together are responsible for the signals from different periphery receptors. According to the classical model of somatosensory processing from NHPs studies (Delhaye et al., 2018; Mountcastle, 2005), the local features such as roughness are processed by the cutaneous receptors, which are conveyed to area 3b, while the global features such as shape are handled by proprioceptive receptors, which project to area 3a. Then, neural signals from areas 3a and 3b project to areas 1 and 2, where the cutaneous and proprioceptive information are integrated. Evidence from a recent study (Kim et al., 2015), however, challenges this prevalent model by finding that area 3b also responds to both cutaneous and proprioceptive inputs. These findings imply that the haptic assessment of 3D objects requires the integration of cutaneous and proprioceptive inputs at all four sub-regions of S1, and these initial processing steps are thought to shape the basic features of the object. Compared to S1, M1 (area 4) is more likely accountable for kinesthetic information processing, such as hand motion and finger positions during the haptic exploration (Gurtubay-Antolin et al., 2018; Kassuba et al., 2013; Masson et al., 2016; Sathian et al., 2011).

Apart from the PSC including S1 and M1, recent human neuroimaging studies of a variety of tasks have observed somatosensory responses in multiple higher-level regions (Sathian, 2016). Specifically, the parietal opercular cortex has been defined as secondary somatosensory cortex (S2) (Burton, 1986), which responds to all types of sensorimotor inputs such as object shape, size, and roughness. Further, S2 is known to bidirectionally connect to the posterior parietal cortex (PPC), to the prefrontal cortex (PFC) and to the premotor cortex (PMC) during both haptic and tactile object processing (Eickhoff et al., 2010, 2008; Rajaei et al., 2018; Sathian et al., 2011; Yang et al., 2017, 2014; Yu et al., 2018b). Although the precise contributions of each area have not yet been established, the sub-regions, including the anterior part of the superior parietal lobule (SPL, areas 5 and 7) and inferior parietal lobule (IPL, area 40) of the PPC, have long been associated with object local and global features processing itself (Sathian, 2016). In contrast, the intraparietal sulcus (IPS) is strongly connected to the bilateral PFC and PMC, both of which have been implicated in planning complex cognitive behavior, attention, decision making, etc. (Hunt et al., 2018; Nee and D’Esposito, 2016; Tremel and Wheeler, 2015). However, it remains unclear how the PSC interacts with these higher-level regions to process haptic information across the cortical hierarchy.

The aim of the present functional magnetic resonance imaging (fMRI) study is to investigate the cortical processing underlying haptic 3D object perception. To manipulate object local and global properties, we designed a series of unique haptic stimuli set combined object local features (roughness) and global features (3D curve), which are changed in a parametric manner. During the fMRI scan participants explored one of sixteen curved surfaces having different roughness (4 kinds of curve × 4 kinds of roughness) in 5-sec event blocks, and were told to estimate one of the two features dependent on the task instruction (i.e., how many curves or how rough the surface was). This experimental design combined with the stimuli set allowed us: (1) to isolate and compare regions across the whole brain relative to surface curve and/or roughness estimation; (2) to test whether the brain regions show parametric variation based on each surface feature; (3) to observe the brain activity across exploration phase and to reveal the interaction between PSC and other higher-level regions as a function of time for each surface feature.

## 2. Materials and methods

### 2.1 Participants

Twenty healthy right-handed volunteers (10 males and 10 non-pregnant females; age range 20-30 years, with mean 22 ± 0.63 years) participated in the fMRI experiments. None of the participants reported a loss of tactile sensation, a history of major medical or neurological illness, such as epilepsy, significant head trauma, or history of alcohol dependence. All of the participants gave written informed consent under an NIH Combined Neuroscience Institutional Review Board-approved protocol (93-M-0170, ClinicalTrials.gov identifier: NCT00001360) in accordance with the Belmont Report and US Federal Regulations that protect human participants.

### 2.2 Finger somatotopic mapping run

One of our research questions was to investigate how the haptic object estimation modulates the activity in the contralateral S1. Thus, to select precise finger areas in the contralateral S1, we first performed a somatotopic mapping run for the right four fingers (index, middle, ring and pinky) using an on-off block design. The duration of each on-phase (stimulation) was 17.5-sec, followed by a 10.5-sec or 14-sec duration off-phase (with off-duration randomly chosen). This on/off-phase cycle was repeated five times for each finger (a total of twenty cycles). The experimenter stands at the entrance of the scanner bore to apply the finger stimulation. During the on-phase, each of the four fingers were randomly and independently poked at a frequency of 4-5 Hz. The participants were instructed to keep their attention on the poked fingertip during the on-phase.

### 2.3 Haptic roughness and curve estimation task run

#### 2.3.1 Haptic stimuli

A total of 17 kinds of 3D printed haptic stimuli were used in the present study. **Figure 1a-d** shows the detailed parameters of the haptic stimuli. Specifically, **Figure 1a** shows four kinds of global curved surfaces, which consist of 1, 2, 3, or 4 curves. **Figure 1b** shows four kinds of local textured surface, consisting of tetragonal arrays of hemispheroidal raised dots with an identical distance center-to-center between adjacent dots in each row: 2, 3, 4, or 5 mm. The hemispheroidal dots had 1 mm diameter and were raised by 1.5 mm from the surface (**Figure 1c**,). All four types of dot patterns were printed on four different curved surfaces with a 40 × 100 mm^2^ rectangular base (**Figure 1d**). **Figure 1e** shows the example of four stimuli with dot spacing equal 5 mm. In total, there were sixteen haptic stimuli (4 kinds of curve × 4 kinds of roughness) for the curve and roughness estimation tasks (**Figure 1f**). Furthermore, to control the basic sensory input by the finger-surface contacts and finger motion, one flat surface without dots was used in the visual motion control task (**Figure 1g**). Three custom-designed, metal-free stimuli containers were used to present all stimuli in a pseudo-random order to the participants during the fMRI experiment. All stimuli shifting occurred during the pre-trial interval, which was manually controlled by the experimenter standing by the MRI bore.

**Figure 1.**
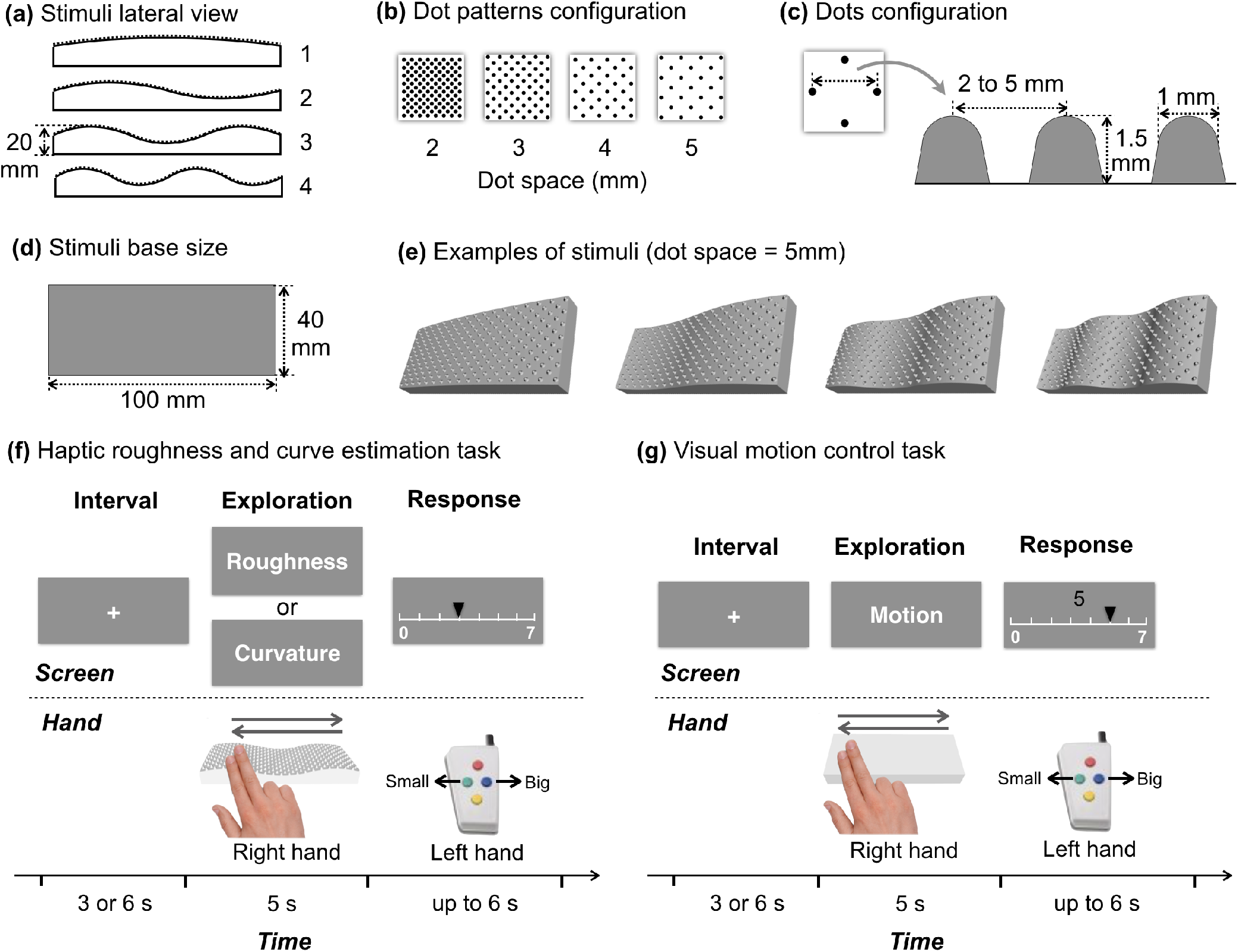
Physical characteristics of the haptic stimuli and the experimental tasks. (a) Lateral view of four kinds of curved stimuli. (b) Four kinds of haptic surfaces consisted of tetragonal arrays of dots with identical dot spacing. (c) Dots heights are 1.5 mm and dot diameters are 1 mm. (d) All four kinds of dot patterns were printed on four different surfaces with a 40 × 100 mm rectangular base. (e) Examples of the stimuli with dot spacing equal to 5 mm. (f) An example trial of roughness and curve estimation task. First, participants were asked to fixate on the visual screen. After a short interval, one of three visual instruction was presented on the screen for 5 s. During this time, participants were asked to move their right index and middle fingers right-and-left along with the surface twice in 5 s and to perform different task based on the instruction: “**Roughness**” estimated the roughness of the surface; “**Curvature**” estimated the number of curves of the stimuli. Then, the participants were asked to assign a number (1 to 7) to the roughness or curves using the button box in their left hand. (g) The visual motion control task used the same procedure, but a flat smooth surface was presented and the participants were asked to just move their fingers right-and-left along with a flat smooth surface twice. At the end of each trail, the participants were asked to move the triangle to the numeric location showed on the center of the screen.

#### 2.3.2 Procedures

Each participant performed four fMRI task runs targeting roughness estimation (RE) and curve estimation (CE). First, to ensure the participants move their fingers at a constant speed, they performed 10-20 trials outside of the scanner. This training will be finished by the experimenter until the experimenter confirmed the uniform motion. The duration of each task run was about 11 min. Participants were informed that a series of surfaces would be presented. Their task was to estimate the roughness or curve of each stimulus, as directed by instructions on the screen (BOLDscreen, Cambridge Research). Roughness was not defined for the participants; instead they were specifically asked to use their own personal definition of haptic roughness. The participants were instructed to choose a comfortable contact force, and to use the same contact force across all trials. The participants’ estimation scale was then established by presenting the smoothest and roughest stimuli. Participants were told that these were two illustrative examples. Participants were asked to assign a whole number from 1 to 7 that seemed appropriate to each surface.

At the start of each task run, the participants were asked to fixate on the center of the screen. After a short interval (3-sec or 6-sec), one of three visual instructions was presented on the screen for 5 sec (see **Figure 1f**). The participants were then asked to move their right index and middle fingers in a right-and-left motion along the surface twice during a 5-sec period at a constant speed and then response to the different stimuli feature task depending on the instruction. When the word “***Roughness***” was presented, the participants were asked to ignore the curve of the stimuli and to estimate the roughness of the surface. In contrast, when the word “***Curvature***” was presented, the participants were asked to ignore the roughness of the surface and to estimate the number of curves. Then, the participants assigned a number to the roughness or number of curves using the button box in their left hand during the 6-sec response phase.

In the visual motion control (VMC) task, we used the same procedure as RE and CE task, while a flat smooth surface was presented (see **Figure 1g**). The participants were asked to just move their fingers in a right-and-left motion along with a flat smooth surface twice. Then, the participants were asked to move the triangle to the numeric location shown on the center of the screen during the 6-sec response phase. These trials were later used as a control for the visual stimuli, and the motor components of the right-hand exploration for the roughness/curvature estimation and the left-hand rating scale selection.

### 2.4 Image acquisition

MRI scans were performed on each participant using a GE Discovery MR750 3T MRI scanner (GE Healthcare, Chicago, IL). No participant was in the scanner for longer than 120 minutes per session. Each scanning session consisted of acquiring a set of fMRI datasets: first, an individual finger somatotopic mapping run of 10 min (240 volumes) duration, and then four haptic task runs each of 11 min duration (265 volumes). Due to the time limitations, 3 out of 20 participants performed only three haptic task runs. Standard T2*-weighted echo planar imaging (EPI) sequence parameters were used to obtain the functional images and ten reverse-blip volumes as follows: repetition time (TR) = 2500 ms; echo time (TE) = 30 ms; phase encoding = A to P; flip angle = 75°; matrix = 77 × 77; 42 axial slices; inplane field of view: 186 × 186 mm^2^; in-plane resolution: 2.58 × 2.58 mm^2^, and 3.0 mm slice thickness (whole brain coverage). After the fMRI acquisition, a T1-weighted magnetization prepared rapid gradient echo (MPRAGE) high-resolution anatomical volume was obtained: voxel size = 1.0 × 1.0 × 1.0 mm^3^, TR = 7040 ms, TE = 3480 ms, matrix = 256 × 256 × 172, duration = 5 min.

### 2.5 Behavioral data analysis

The RE and CE estimates (scale values) and response times of each participant (see **Figure 2**) were collected with open-source application PsychoPy software v1.85.0 (Peirce et al., 2019). The R programming language (R Core Team, 2013) was used for the additional statistical analyses.

**Figure 2.**
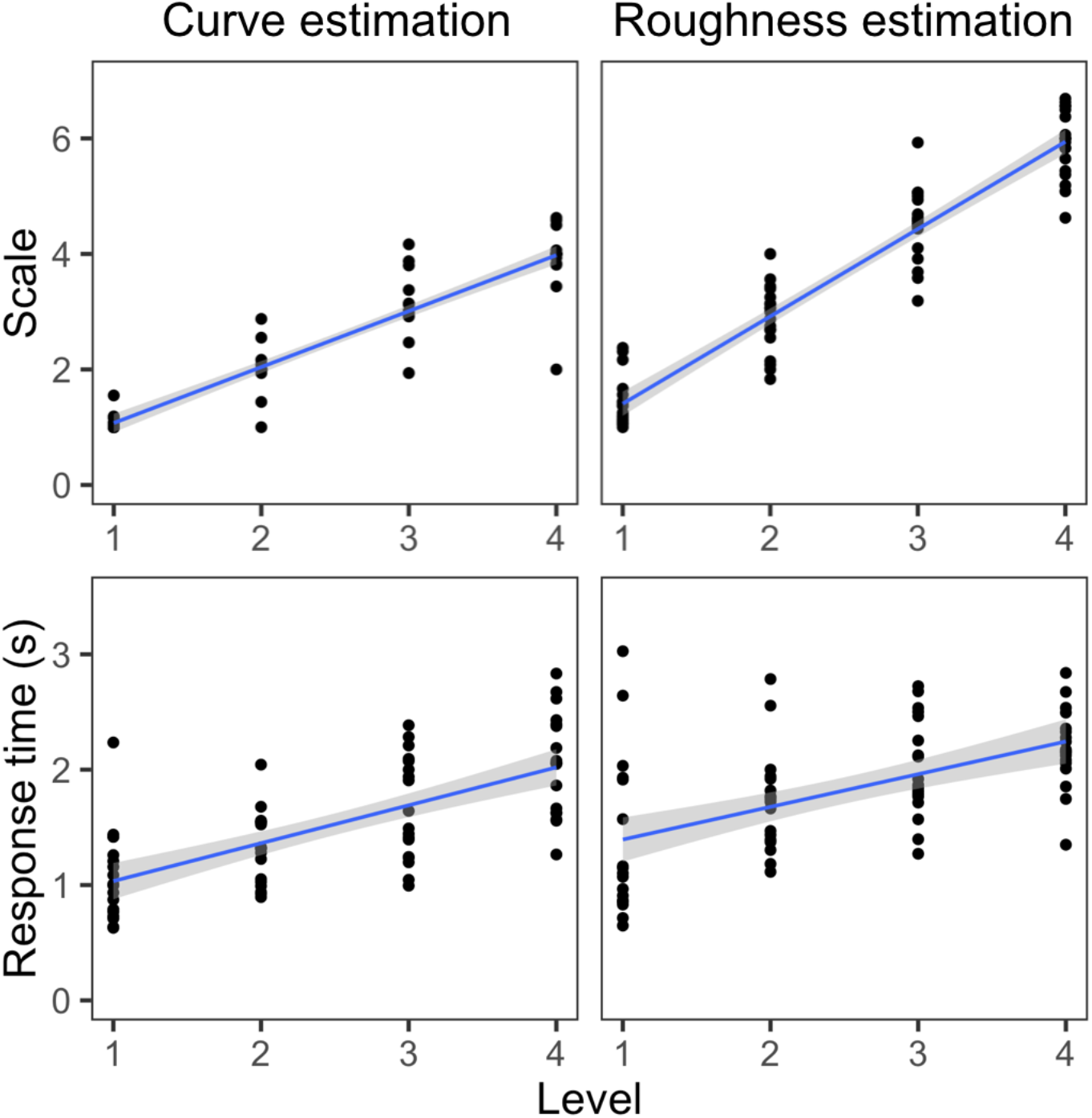
Behavioral performance of curve and roughness estimation tasks. Black dots represent the average scale or response time of each participants. The blue lines represent the linear regression line. The gray back ground represents the 95% confidence interval.

### 2.6 fMRI data analyses

FMRI data were analyzed using “*afni_proc.py*” with the AFNI/SUMA (version = 18.1.08) software package (http://afni.nimh.nih.gov/) (Cox, 1996; Saad et al., 2006). Cortical surfaces for each participant were created using FreeSurfer (Version 6.0) (http://surfer.nmr.mgh.harvard.edu/) (Fischl, 2012) by running “*recon-all*” on each T1-weighted anatomical and converting the results to standard NIFTI/GIFTI format with AFNI’s “*@SUMA_MakeSpec_FS*”.

#### 2.6.1 Individual participant: preprocessing and modeling

The full “*afni_proc.py*” command used to generate the processing stream and quality control is provided in the ***Supplementary material***. We briefly describe the implemented processing blocks and options used here. Before statistical analysis, the first two volumes of each run were removed, and slice-timing correction was then performed to adjust for differences in slice-acquisition times. Then, we applied blip up/down non-linear alignment to all EPI images and then aligned all images to participants’ own anatomical image (Glen et al., 2020). Motion correction with rigid body (three translation and three rotation) alignment was performed, and volumes with Euclidean norm (enorm) of the rigid body motion parameters greater than 0.3 mm were censored. EPIs were mapped to the surface domain and blurred to a smoothness of 6 mm FWHM on the surface. Finally, each node’s time series was scaled mean of 100, so that time series fluctuations would correspond to interpretable units of local blood-oxygen level dependent (BOLD) percent signal change (Chen et al., 2017).

Within the “*afni_proc.py*” command, a general linear model (GLM) was also fitted to the fMRI data for each participant. The BOLD signal was modeled for the finger somatotopic mapping run and all haptic RE and CE task runs with block function convolved with the canonical hemodynamic response function (HRF) using AFNI’s “*3dDeconvolve*”. Assuming a first-order autoregressive model, the serial autocorrelation was estimated from the pooled active nodes with the restricted maximum likelihood procedure. The motion-related artifacts were minimized via the incorporation of six parameters (three translation and three rotations) from the rigid-body realignment stage into each model. The estimates were evaluated using linear contrasts of finger relative to baseline in each participant or each task. Furthermore, aside from the visual data confirmation, the output of AFNI’s “*@ss_review_basic*” for each single participant processing was used for quality control, which contains the max motion, tSNR, smoothing values, counts of outliers, etc.

Then, we obtained the sub-brick images for (section 2.6.2) localizing the mean specific S1 sub-regions, (section 2.6.3) observing the whole-brain activity pattern of CE and RE tasks, (section 2.6.4) observing the brain regions parametrically modulated by CE and RE tasks. Finally, we also performed (section 2.6.5) a region of interest (ROI) analysis to observe the time series data from the contralateral S1 and other high-level regions.

#### 2.6.2 Group analysis: localize specific S1 sub-regions for index and middle fingers

First, an one-sample t-test was used to confirm the activation of each finger (index, middle, ring and pinky) from the finger somatotopic mapping run. The node-wise significance threshold was set at p < 0.005 (t_19_ > 3.17, two-sided testing) (Chen et al., 2019). Then, we classified activations for index and middle fingers around the postcentral gyrus (poCG) into four sub-regions (area 3a, 3b, 1, 2) within S1 and M1 (area 4). There are several anatomical landmarks for identifying the sub-regions of human S1. Specifically, area 3a and 3b are known to locate along the posterior bank of the central sulcus (CS), area 1 occupies the crown of poCG, and area 2 lies on the posterior bank of poCG. Furthermore, hand area 4 is known to locate on the hand knob of the preCG. Based on these landmarks, we generated the masks of area 4, 3a, 3b, 1 and 2 according to the averaged activations of index and middle fingers (**Figure 3a-b**).

**Figure 3.**
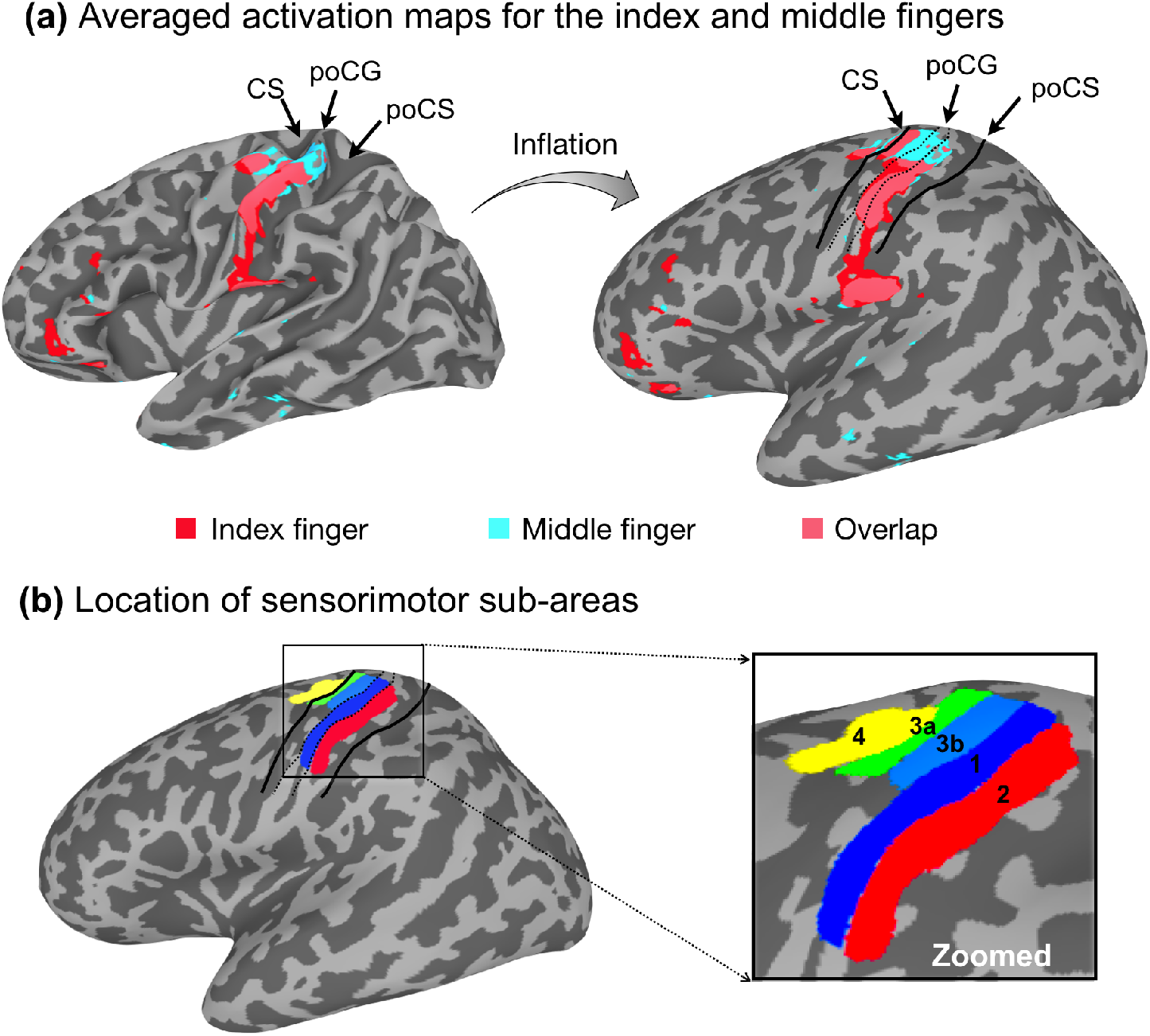
Somatotopic mapping for right index and middle fingers. (a) Illustrated the averaged activation maps for the index and middle fingers. For all participants, finger maps were observed within the CS, poCG and poCS contralateral to the stimulated fingers. Further, we found that the middle finger was localized to a more superior position than index finger. (b) Illustrated the location of sensorimotor sub-areas. Based on the landmarks, we generated the masks of area 4, 3a, 3b, 1 and 2 according to the averaged activations of index and middle fingers. CS, central sulcus; poCG, postcentral gyrus; poCS, postcentral sulcus.

#### 2.6.3 Group analysis: average activity modulation by CE and RE tasks

First, an analysis of variance (ANOVA) was used to confirm the whole brain activation of each task (RE, CE, and VMC). We then evaluated the contrast of mean of the RE task with the mean of the VMC task (RE – VMC) [**Figure 4a**] and the contrast of mean of the CE task with the mean of the VMC task (CE – VMC) [**Figure 4b**]. We then evaluated the contrasts of (CE – VMC) – (RE – VMC) [**Figure 4c**] and (RE – VMC) – (CE – VMC) [**Figure 4d**] in order to identify brain regions affected by the haptic curve and roughness estimation. The height threshold was set at p < 0.005 (t_19_ > 3.17, two-sided testing). The statistical threshold for the spatial extent test on the nodes over the whole brain was set at p < 0.05, the minimum significant clusters was 266 mm^2^.

**Figure 4.**
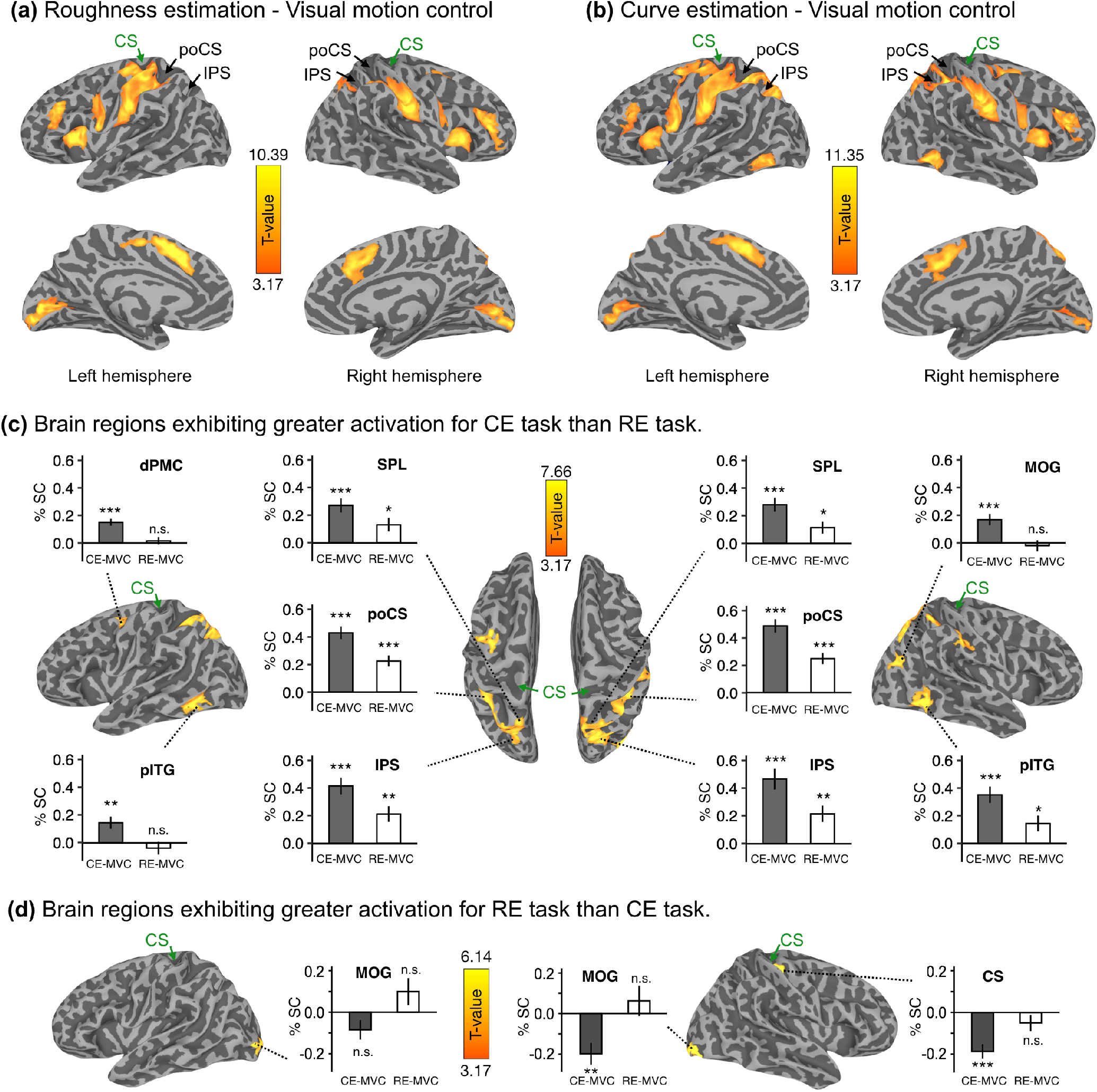
Mean (n=20) brain activation of roughness estimation (RE) and curve estimation (CE) rendered on the cortical surfaces using SUMA. (a) Brain regions exhibiting greater activation for RE task than visual motion control (VMC). (b) Brain regions exhibiting greater activation for CE task than VMC. (c) Brain regions for (CE - VMC) vs. (RE - VMC) contrast. (d) Brain regions for (RE - VMC) vs. (CE - VMC) contrast. The extent threshold of activation was p < 0.05 (area larger than 266 mm^2^) corrected for each search node with the height threshold of p < 0.005 (t value larger than 3.17) uncorrected. These bar graphs represented the mean activation in each ROIs for each task (n=20). The error bars indicate the standard error of the mean (SEM). CS, central sulcus; poCS, postcentral sulcus; IPS, intraparietal sulcus; dPMC, dorsal premotor cortex; SPL, superior parietal lobule; pITG, posterior part of the inferior temporal gyrus; MOG, middle occipital gyrus. Asterisks represented the statistically significant of one-sample t-test. *: p < 0.05, **: p < 0.01, ***: p < 0.001, n.s.: not significant.

#### 2.6.4 Group analysis: parametric main effects of CE and RE tasks

To locate any regions that showed a parametric response to CE and RE, we next performed a whole brain group analysis with parametric regressors (Chen et al., 2014). We identified areas exhibiting activation that was positively or negatively associated with curve (**Figure 5a**) and roughness (**Figure 5b**). Parametric predictors for the time course were modeled as a boxcar function for each condition, with the amplitude equal to the roughness and curve levels. Each type of stimulus has been set at four levels in the model, which (−3, −1, 1, 3) normalized to have a zero-mean, and subsequently convolved with the HRF. Both of the height threshold and spatial extent thresholds were set to the same as above.

**Figure 5.**
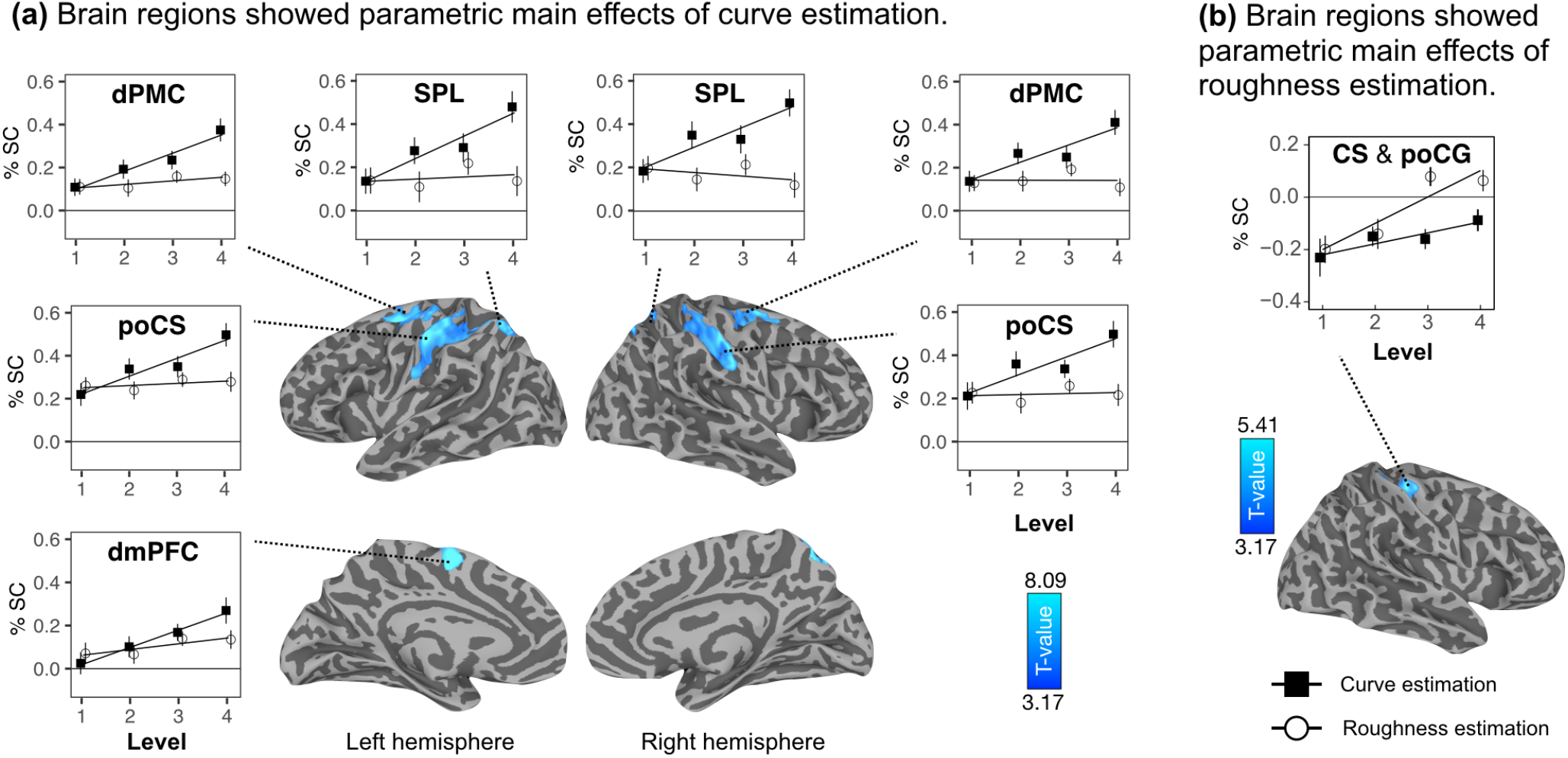
Brain regions showed parametric main effects of (a) curve estimation and (b) roughness estimation. These plots represented the mean activation in each ROIs for each task per stimuli level across participants (n=20). For curve estimation task, level 1 to 4 represent number of curves. For roughness estimation task, level 1 to 4 represent the surface roughness. The error bars indicate the standard error of the mean (SEM). The extent threshold of activation was p < 0.05 (area larger than 266 mm^2^) corrected for each search node with the height threshold of p < 0.005 (t value larger than 3.17) uncorrected. CS, central sulcus: poCG, postcentral gyrus; poCS, postcentral sulcus; SPL, superior parietal lobule; dPMC, dorsal premotor cortex; dmPFC, dorsomedial prefrontal cortex.

#### 2.6.5 Region of interest (ROI) analysis

After the whole brain analysis, we extracted the time series data from the following ROIs to examine activation at each time point during RE and CE tasks. To examine the activation in contralateral PSC, we first defined contralateral area 4, 3a, 3b, 1 and 2 as mentioned above. Furthermore, to examine the activation in curve or roughness sensitive areas, we defined the ROIs that showed a parametric response to curve estimation (**Figure 5a**). Then, we extracted all the functional time series signals at these specified ROIs for all participants from the scaled EPIs in the surface space. Finally, these signals were convolved with a gamma variate HRF (default gamma HRF of 3dDeconvolve in AFNI) to produce estimate the response amplitude (% signal change) for each ROI to each task trial. The averaged activation profiles of each ROI are summarized in **Figure 6 and 7**.

**Figure 6.**
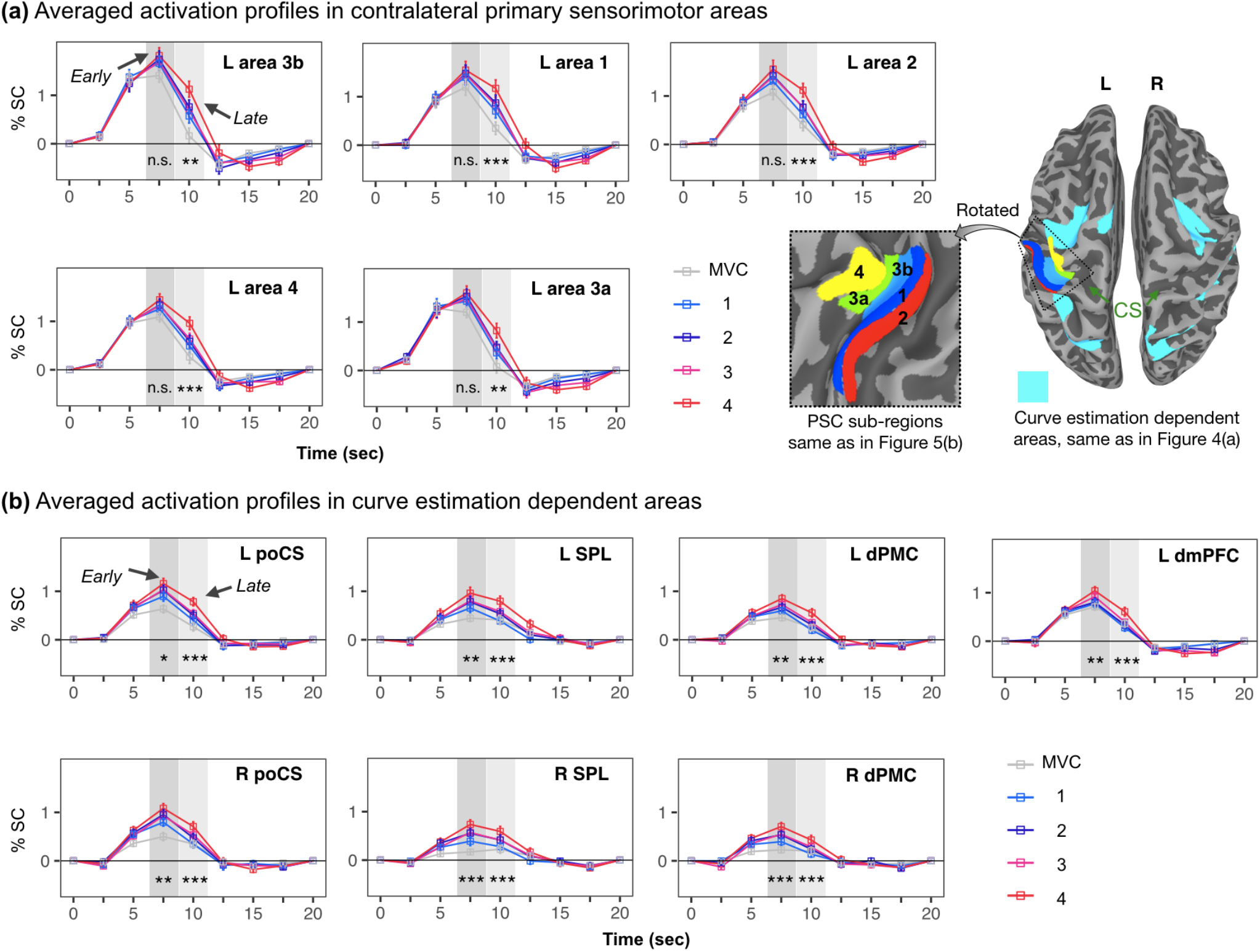
Activation profiles at each time point during haptic curve estimation. (a) Averaged activation profiles in contralateral sensorimotor areas. (b) Averaged activation profiles in areas sensitive to curve estimation as shown in Figure 3(a). The error bars indicate the standard error of the mean (SEM). The darker gray square in each plot represented the activation peak for the early exploration phase and light gray square represented the activation peak for the late exploration phase.

**Figure 7.**
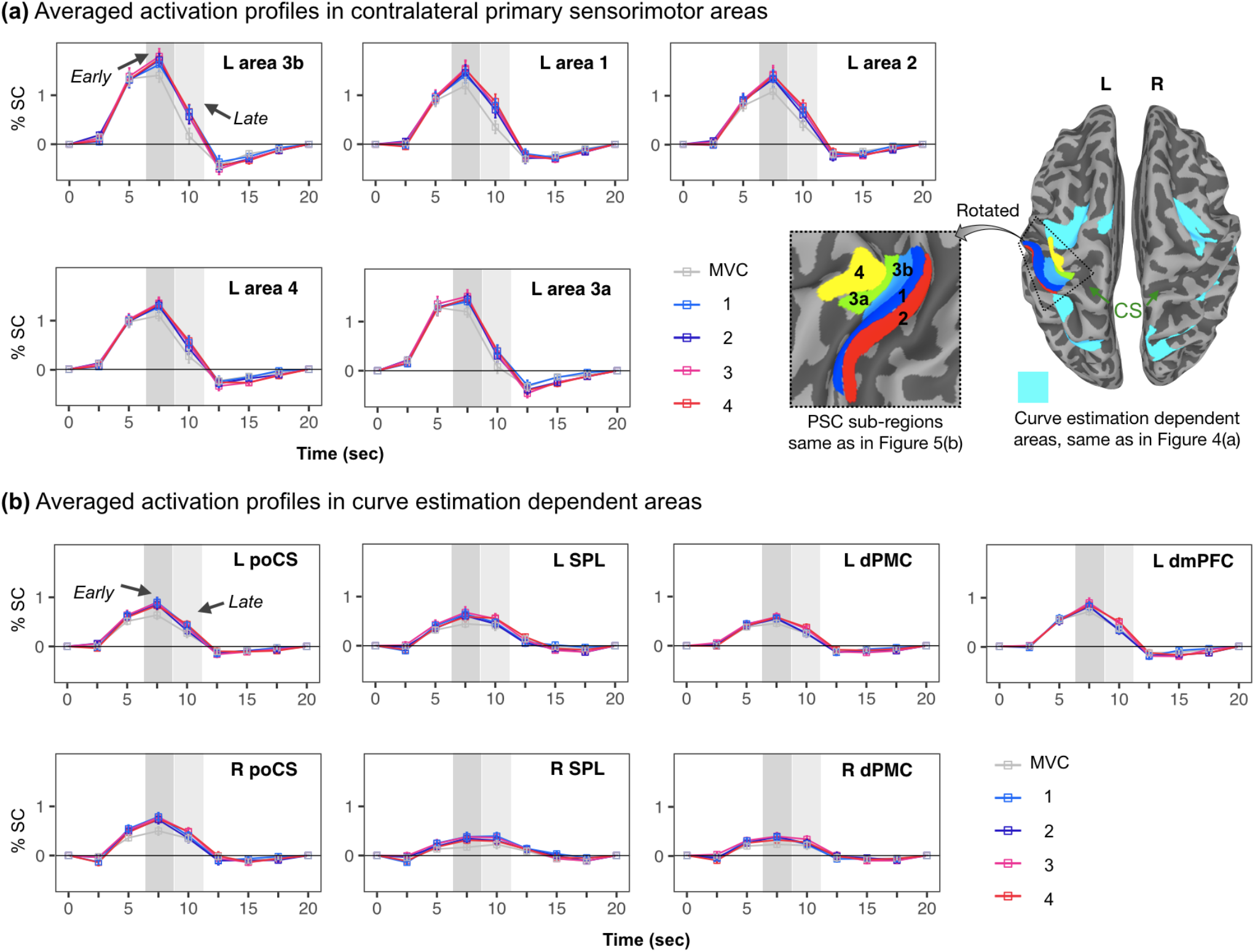
Activation profiles at each time point during haptic roughness estimation. (a) Averaged activation profiles in contralateral sensorimotor areas. (b) Averaged activation profiles in areas sensitive to curve estimation as shown in Figure 3(a). The error bars indicate the standard error of the mean (SEM). The darker gray square in each plot represented the activation peak for the early exploration phase and light gray square represented the activation peak for the late exploration phase.

## 3. Results

### 3.1 Behavioral performance

Eighteen of twenty participants (For technical reasons, behavioral data for two participants was lost) were included in the behavioral data analysis. As shown in **Figure 2**, all participants were able to scale the roughness and curve of the stimuli. The linear regression analysis (blue lines in **Figure 2**) revealed that the scale values of both roughness [r^2^ = 0.744, p < 0.001] and curve [r^2^ = 0.875, p < 0.001] estimation were significantly increased dependent on the stimuli level. In addition, we also performed the linear regression analysis on the mean response times for all tasks (relative to the offset of the stimulus). We found that the response times were significantly increased dependent on the stimuli level in both roughness [r^2^ = 0.302, p < 0.001] and curve [r^2^ = 0.465, p < 0.001] estimation tasks. Furthermore, a two-way (two tasks × four stimuli levels) repeated measures ANOVA of the mean response time also revealed significant main effects of task [F_1, 17_ = 14.38; P = 0.001] and stimuli level [F_3, 51_ = 43.58; P < 0.001], but no significant interaction between task and stimuli level [F_3, 51_ = 1.49; P = 0.228]. In detail, the multiple comparisons with Bonferroni correction revealed that the difference between RE and CE tasks at level two is significantly (difference is 0.4±0.11 sec, P = 0.020) but not for all other levels (Ps > 0.05). These results suggest that haptic estimation of curves and roughness are comparable.

### 3.2 fMRI results

#### 3.2.1 Somatotopic mapping for right index and middle fingers

For all participants, finger maps were observed within the CS, poCG and poCS contralateral to the stimulated fingers. **Figure 3a** shows the averaged (n = 20) activation maps for the index (red) and middle (water blue) fingers. In consistent with the previous studies (Ann Stringer et al., 2014; Martuzzi et al., 2014), we found that the middle finger was localized to a more superior position than index finger.

Then, we defined index and middle finger corresponded sensorimotor sub-regions relative on the anatomical landmarks on CS, poCG and poCS (**Figure 3b**).

#### 3.2.2 Whole brain activation for roughness and curve estimation

Initially, we confirmed that both of the RE and CE tasks (relative to VMC) activated a widespread set of brain regions including contralateral CS, preCG and poCG (**Figure 4ab**). In addition, we also found significant activations on bilateral ventrolateral prefrontal cortex (vlPFC), dorsal and ventral premotor cortex (dPMC, vPMC), insular cortex, parietal operculum (PO), dorsomedial prefrontal cortex (dmPFC) and calcarine sulcus, and right poCS, right poCG, right intraparietal sulcus (IPS), right superior parietal lobule (SPL). Moreover, we also found that CE task, but not the RE task, significantly activated bilateral posterior part of the inferior temporal gyrus (pITG), and left IPS, left SPL.

#### 3.2.3 Whole brain activation for curve estimation stronger than roughness estimation and vice versa

As shown in **Figure 4c**, the regions including bilateral poCS, IPS, SPL and pITG, as well as left dPMC, and right MOG were activated stronger for CE than RE task. In contrast, the contrast of (RE – VMC) – (CE – VMC) only showed stronger activation in bilateral MOG and right CS (**Figure 4d**).

#### 3.2.4 Brain regions showed parametric main effects of roughness and curve estimation

**Figure 5a** shows the activation of these brain regions linear increased as the number of curves were increased. These include bilateral dPMC, poCS, SPL, and left dmPFC. Then, we extracted the mean activity signal (% signal change) of each of these seven clusters for CE and RE tasks, to confirm the linear relationship between stimuli level and brain activation. As shown in the plots of **Figure 5a**, the brain activations of these regions showed linear increases relative to the curve estimation level (black squares), but not for roughness estimation level (outline circles). Compared to curve estimation, we uniquely identified right CS extended to poCG (CS & poCG) were parametrically modulated by the roughness estimation (**Figure 5b**).

#### 3.2.5 ROI results for roughness and curve estimation

The averaged fMRI time series data in five contralateral PSC sub-regions and seven curve dependent higher-level regions were shown in **Figure 6** and **7**. Note, since the anterior part of left poCS as shown in **Figure 5a** was overlapped with the area 2 as defined by the somatotopic mapping (**Figure 3b**), we used un-overlapped posterior part of poCS here to extract the time series data in **Figure 6** and **7**. A linear mixed-effects model analysis was performed using the “*lme4*” package in R (Bates et al., 2014) to evaluate the brain activity time series of each ROI. Here, we set time factors at two levels, which corresponding to the early (time point of 7.5 sec in **Figure 6** and **7**) and late (time point of 10 sec) part of the stimuli exploration phase. The stimuli factors were set at four, corresponding to stimuli curves or roughness. If there was a significant interaction between time and stimuli level, a post hoc test was conducted for the simple effect of time. These p values were Bonferroni-corrected. Surprisingly, we found that contralateral PSC sub-regions showed significantly different activation patterns from those found in higher-level regions (**Figure 6ab**). Specifically, for contralateral PSC sub-regions (**Figure 6a**), we only found the parametric modulation by curve estimation during the late phase, but not for the early phase. In contrast, the higher-level regions, for example the activation of left poCS (**Figure 6b**), we found a linear increase relative to the curve estimation level both for the early and late phase. In contrast, as shown in **Figure 7ab**, we did not find the similar time dependent activation profiles nether in the contralateral PSC sub-regions nor higher-level regions for the roughness estimation.

## 4. Discussion

In the present study, we investigated the brain activity during haptic curve and roughness estimation using a parametric fMRI experiment. Our results extend beyond the previous findings (Mueller et al., 2019; Sathian et al., 2011; Stilla and Sathian, 2008) by revealing brain regions that show parametric variation that is dependent on the curve numbers (**Figure 5a**). Further, our experimental design enabled us to observe the brain activity across exploration phase (i.e., early and late phase) in contralateral PSC and other higher-level regions during the haptic curve and roughness estimations. In particular, we found that only activation of the higher-level regions showed linear increases relative to the number of curves through the exploration phase (**Figure 6b**), whereas we found that the contralateral PSC (**Figure 6a**) were parametric modulated by the number of curves only during the late exploration phase. In contrast, these time-dependent brain activity features in the same sort of regions did not appear during the haptic roughness estimation (**Figure 7**). Together, our findings suggest that haptic 3D object perception is processed across the cortical hierarchy, while the contralateral PSC interacts with other higher-level regions across time in a manner that is dependent upon object features.

First, we confirmed that haptic curve estimation enhanced a sort of higher-level regions (**Figure 4c**) than that of haptic roughness estimation which has been well discussed in the previous studies (Mueller et al., 2019; Sathian et al., 2011; Stilla and Sathian, 2008). Haptic perception usually reflects the sensory processing from cutaneous receptors in the skin and proprioceptive receptors in the muscles, joints, and so on (Lederman and Klatzky, 2009; Sathian, 2016). In the present study, participants were asked to use the same approach to explore the same stimuli in both CE and RE tasks while estimate one of each surface features follows the instruction. This approach has the advantage of keeping the stimuli constant across tasks, as well as both curve and roughness estimation would be determined by the similar amounts of cutaneous and proprioceptive inputs. Further, since we found that all participants can scale both curve and roughness properly in a few seconds (**Figure 2**), we can assume that attentional demands were comparable between CE and RE tasks. Therefore, it is reasonable to interpret that why we found very similar brain activity maps for CE and RE tasks (**Figure 4ab**). These regions of (CE – VMC) – (RE – VMC) contrast (**Figure 4c**) were thought to contribute to the extraction of the 3D geometric information (curves) from objects or other higher-level processing such as visual imagery (Deshpande et al., 2010; Kassuba et al., 2013; Lacey et al., 2010).

Interestingly, our results highlighted that haptic curve estimation parametrically modulated the activations depend on the exploration phase within the contralateral PSC (i.e., areas 4, 3a, 3b, 1 and 2) and several higher-level regions. Specifically, the whole-brain parametric analysis revealed that the activations of several higher-level regions, including bilateral poCS, SPL, dPMC, and left dmPFC (**Figure 5a**), are uniquely and linearly increase with increasing curve numbers. Nonetheless, looking beyond the fMRI adaptation effects (Barron et al., 2016; Krekelberg et al., 2006; Larsson and Smith, 2012), the ROI analysis also revealed significant curve parametric modulation in all contralateral PSC sub-regions at the late exploration phase, but not the early exploration phase (**Figure 6a**). Such findings suggest the possibility that the contralateral PSC and these higher-level regions interacted differently across the haptic exploration phase for curve estimation.

Human sensory processing is considered typically within a hierarchical framework, consisting of a series of discrete stages from the primary sensory cortex to the whole brain. In the somatosensory system, the sensorimotor information is first projected initially to the contralateral PSC, encoding basic perceptual dimensions, such as edge, roughness and hand motion (Pleger and Villringer, 2013; Sathian, 2016). Then, following higher stages beyond contralateral PSC, the second-order sensory cortex, such as areas 1 and 2, is known to have bilateral receptive fields (Iwamura, 1998), which are sensitive to hand movement direction and object shape, etc (Sathian, 2016). Aside these somatosensory areas, the caudal part of SPL (i.e., area 7) also appears to be involved in the higher-order processing of sensorimotor information. Furthermore, area 7 was known to functional connect with bilateral SPL, dPMC, and dmPFC (putative human supplementary motor area), and this network thought to function in the integrative sensory, motor and cognitive functions (Freedman and Ibos, 2018; Nelson et al., 2010). Therefore, the parametric modulation in these higher-level regions through the exploration phase may be represented the higher-order functions such as curve reconstruction, finger motion control during the exploration, and so on.

In contrast, this strictly bottom-up formation cannot adequately interpret that why we only found the parametric modulation in the contralateral PSC at the late exploration phase, but not at the early exploration phase. One possible interpretation of our finding is related to the bidirectional hierarchy models such as predictive coding theoretical frameworks (de Lange et al., 2018; Friston, 2010), which the lower sensory cortex receives not only bottom-up input but also top-down feedback (Yu et al., 2019b, 2019a, 2018a). Thus, our finding suggest that the prior experience of the curved surface may provide top-down feedback to modulate the contralateral PSC in a parametric manner during the late exploration phase. In the present study, the participants were asked to explore each surface twice during the exploration phase. It is reasonable to assume that the participants would know curve numbers during the first exploration (roughly during the early phase), and the second exploration (roughly during the late phase) was more likely to confirm the answer. Thus, one important insight from these data is that the physical property, such as the curve numbers, might not be parametrically encoded in the contralateral PSC, while the top-down feedback modulation may occur in the contralateral PSC.

It may be surprising that, for roughness estimation, we did not find parametric modulation in the contralateral PSC and these higher-level regions neither for the early nor the late exploration phase (**Figure 7ab**). From the behavioral perspective, this result may reflect that the participants are more likely assign a number to the roughness at the end of the exploration since the roughness is an abstractive definition and may not easy to remember it by a number at the early exploration phase as in the curve estimation case. From the brain function perspective, this finding may reflect the different coding and processing between roughness and curve in the brain. Both the findings, as shown in **Figure 4c** of the present study and previous neuroimaging studies (Kassuba et al., 2013; Stilla and Sathian, 2008), may support this assumption. For example, we found that the activations of bilateral SPL, IPS, and pITG are significantly stronger for curve estimation than those of roughness estimation, and these regions have been demonstrated to tends to be more specialized for visual object processing (Kassuba et al., 2013; Stilla and Sathian, 2008). In contrast, regions such as the S2 are more sensitive to the haptic perception of surface roughness processing (Stilla and Sathian, 2008). Despite this, we cannot exclude the contribution of other factors, such as the basic properties of the roughness stimulus. For example, the previous behavior studies (Dépeault et al., 2009; Eck et al., 2013) have used the dot spacing ranged from 1.5 to 8.5 mm, while in the present study we used the dot spacing ranged from 1 to 5 mm, which limited us to observed the parametric modulation. Much more work is necessary to resolve these issues.

## 5. Conclusion

In summary, even we find that haptic curve and roughness processing share a large proportion of cortical regions, curve estimation parametrically modulated activation in contralateral PSC and bilateral poCS, SPL, dPMC, and left dmPFC, but not for roughness estimation. Further, we find remarkable differences of exploration phase dependent brain activation between the contralateral PSC and higher-level regions related to haptic curve estimation. This finding may represent the nature of time-dependent interaction across the sensory information cortical hierarchy to shape our behavior.

## Acknowledgments

We thank Vinai Roopchansingh and John Andrew Derbyshire for the fMRI sequence used here. We thank Natasha Topolski for support of human volunteer scanning. We thank the NIH machine shop for help with 3d-printing the stimuli and device. The research was conducted as part of the NIMH Intramural Research Program (#ZIA-MH002783).

## Author contributions

J.Y., P.M., Y.Y., and P.B. designed and performed the fMRI experiments. D.H., Y.E., and J.W. contributed to the conception and design. J.Y., P.M., G.C., and P.T. analyzed the fMRI data. J.Y., P.M., Y.Y., and PB wrote the paper. All authors discussed and commented on the manuscript.

## Finding

This work was supported by JSPS KAKENHI Grant Numbers JP20K07722, JP18H01411 and Japan-U.S. Science and Technology Cooperation Program (Brain Research).

## Data and materials availability

All data needed to evaluate the conclusions in the paper are present in the paper. The data presented here are available via an NIH Acronis Access link upon request. Downloading access requires a signed data sharing agreement as part of the intramural IRB protocol.

## Supplementary material

### 1. Individual participant preprocessing and modeling with afni_proc.py

Here we provide the specific processing commands used to generate the full processing stream (through regression modeling) for each participant in this study. The entire pipeline for individuals was created using the afni_proc.py command in AFNI (version = 18.1.08; Cox, 1996), which generates a commented processing script. The running of this script creates a full processing output directory, which contains both intermediate and final datasets for review, and both quantitative and qualitative quality control (QC) features, such as review scripts to browse data and an HTML document showing features of each processing block for efficient review and verification by the research. These features were used to verify the processing in the present study.

**Table S1** provides the main processing stream used to process the curve and roughness estimation runs in the present study. The only input variables here to be specified were subject ID (“subj”) and the top level directory of the group (“top_dir”). The remaining dataset and variable names should mainly be clear by name and context (e.g., the “run*” datasets are the EPI runs, “T1w*” is the T1-weighted anatomical, etc.). We note that the times in the stimulus timing files (*.txt) have already been adjusted to account for the fact that the first 2 TRs of the EPI dsets are removed during processing here.

For potential future studies based on these commands, we note that the following more recent features would be recommended for inclusion. The option “-check_flip” can be added after “align_opts_aea” to help check for EPI-anatomical dataset consistency (for a discussion of left-right flipping issues common in MRI analyses, see (Glen et al. 2020)). The following option can also be added to help check initial data quality and motion removal: “-radial_correlate_blocks tcat volreg”. For improved style of automatic QC HTML document, we recommend adding “-html_review_style python”. Additionally, the argument after “-volreg_align_to” could be changed to “MIN_OUTLIER” instead of “third”, so that the reference volume for motion corrected is selected to be the volume with fewest outliers. Finally, another censoring criterion could be added, which is based on the fraction of outliers in a volume; adding “-regress_censor_outliers 0.05” would censor any volume with more than 5% outliers.

**Table S2** provides the afni_proc.py processing command used to process the task fMRI for somatotopic finger mapping. The same abbreviations, comments and recommendations from the processing script in **Table S2** apply to this command.

**Table S1.**
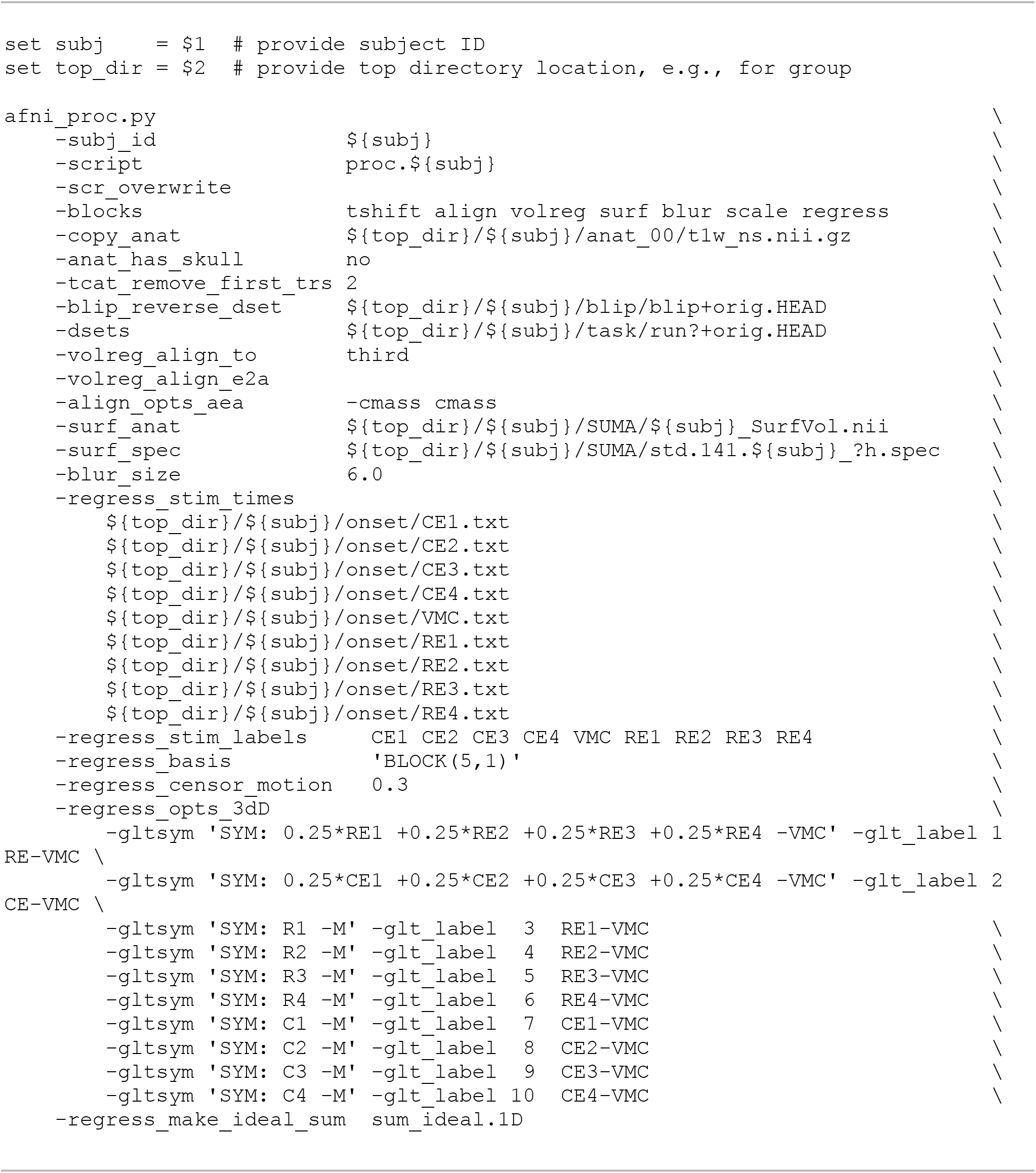
The afni_proc.py command used to process the curve and roughness estimation runs in the present study. In the stimulus timing files and general linear tests (GLTs), the following abbreviations are used to specify task types: CE = curve estimation conditions (4 runs); RE = roughness estimation conditions (4 runs); VMC = visual motion control condition (4 runs). Backslashes are continuation of line characters, allowing for spacing and increased readability. Here and below, tcsh syntax is used to specify variables, loops, etc.

**Table S2.**
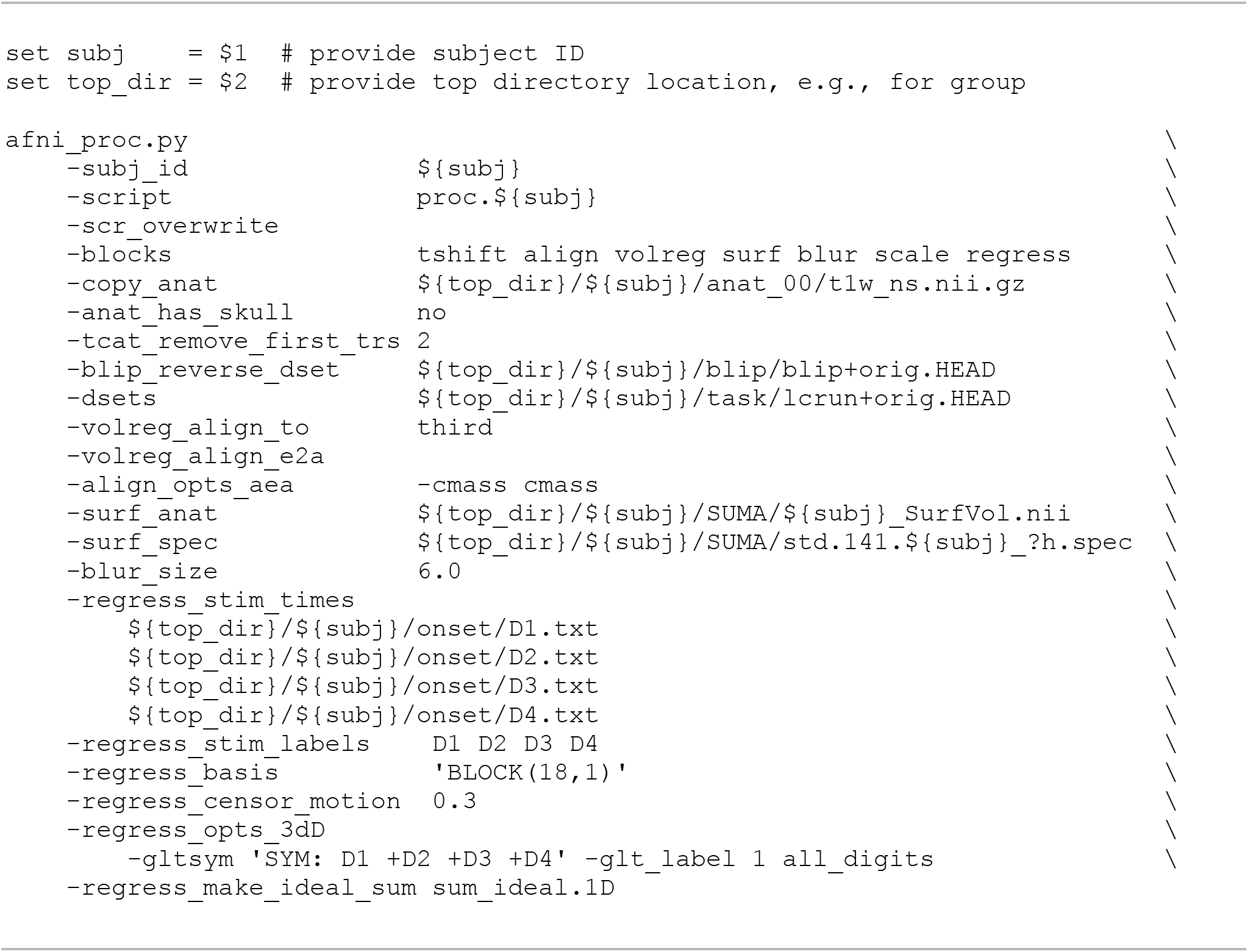
The afni_proc.py command used to process the somatotopic finger mapping runs in the present study. In the stimulus timing files and general linear tests (GLTs), the following abbreviations are used to specify task types: D1 = index finger; D2 = middle finger; D3 = ring finger; D4 = pinky finger.

### 2. AFNI group analysis commands

Here we provide the AFNI commands used to specify and perform the group analysis in this study. In order to simplify the setup of group analysis commands, AFNI contains the gen_group_command.py program to several types of statistical tests, such as t-tests, mixed effects meta analyses (MEMAs), and ANOVAs. This program creates the full command for the relevant statistical program in AFNI, permitting the user to focus on specifying the participants and model formulation rather than programmatic syntax; this simplifies tasks such as adding/removing subjects and varying model specification, as well as reducing bugs in analysis.

**Table S3** specifies the command used to set up a t-test for group analysis to localize specific S1 subregions for the index and middle fingers. **Table S4** specifies the command used to set up an ANOVA to evaluate the contrast of mean of the roughness estimation (RE) and curve estimation (CE) task with the mean of the visual motion control (VMC), as “RE – VMC”. **Table S5** specifies the multivariate modeling (MVM) for AFNI’s 3dMVM (directly, not using gen_group_command.py) for revealing the parametric main effects of curve and roughness estimations. This analysis was performed on the surface, and hence the command is run for both the left and right hemispheres (lh and rh, respectively).

**Table S3.**
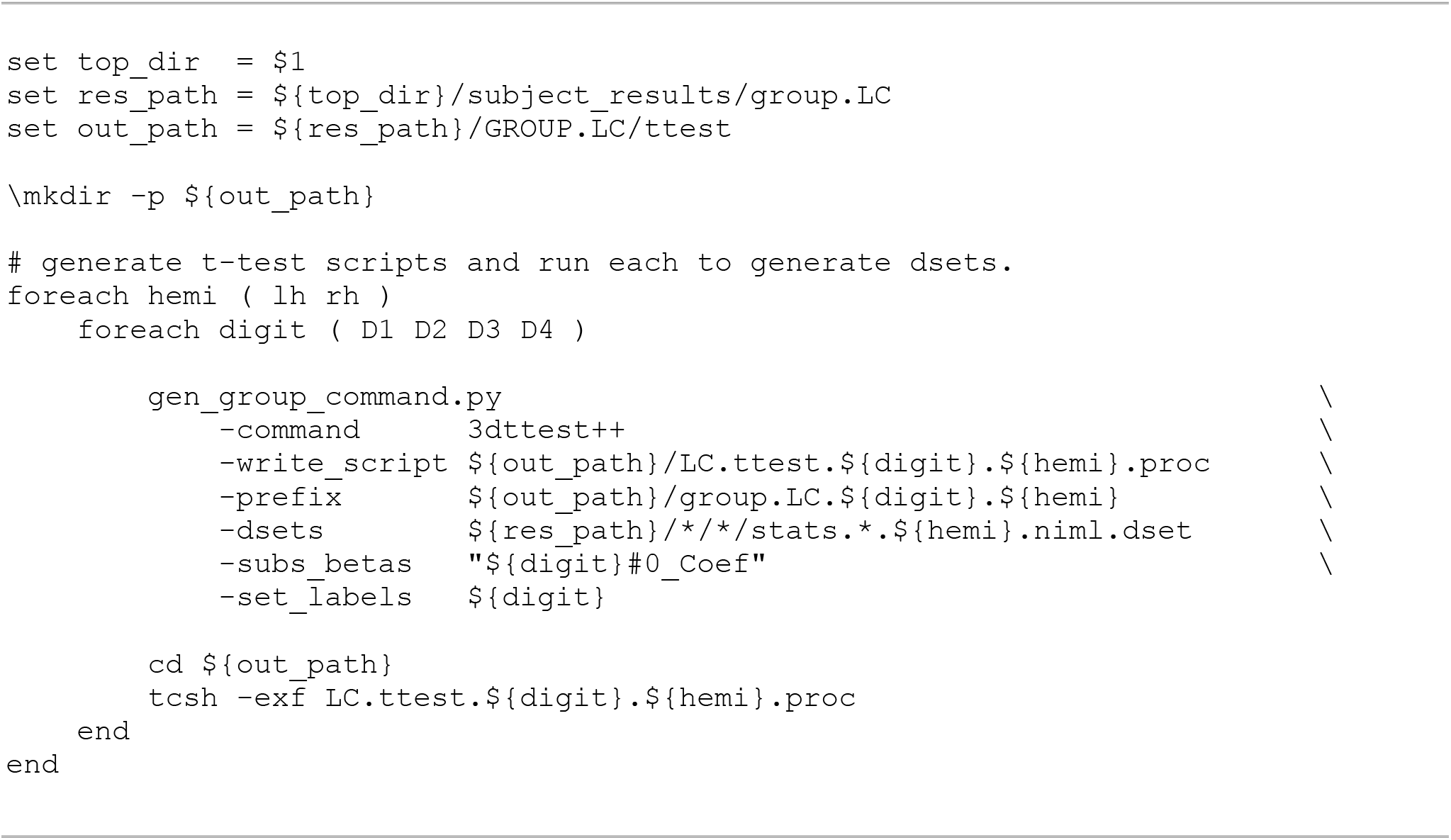
The command to generate t-tests for group analysis to localize specific S1 sub-regions for all four fingers and only the index and middle fingers’ maps were presented in the present study. The following abbreviations are used: lh = left hemisphere; rh = right hemisphere; D1 = index finger; D2 = middle finger; D3 = ring finger; D4 = pinky finger.

**Table S4.**
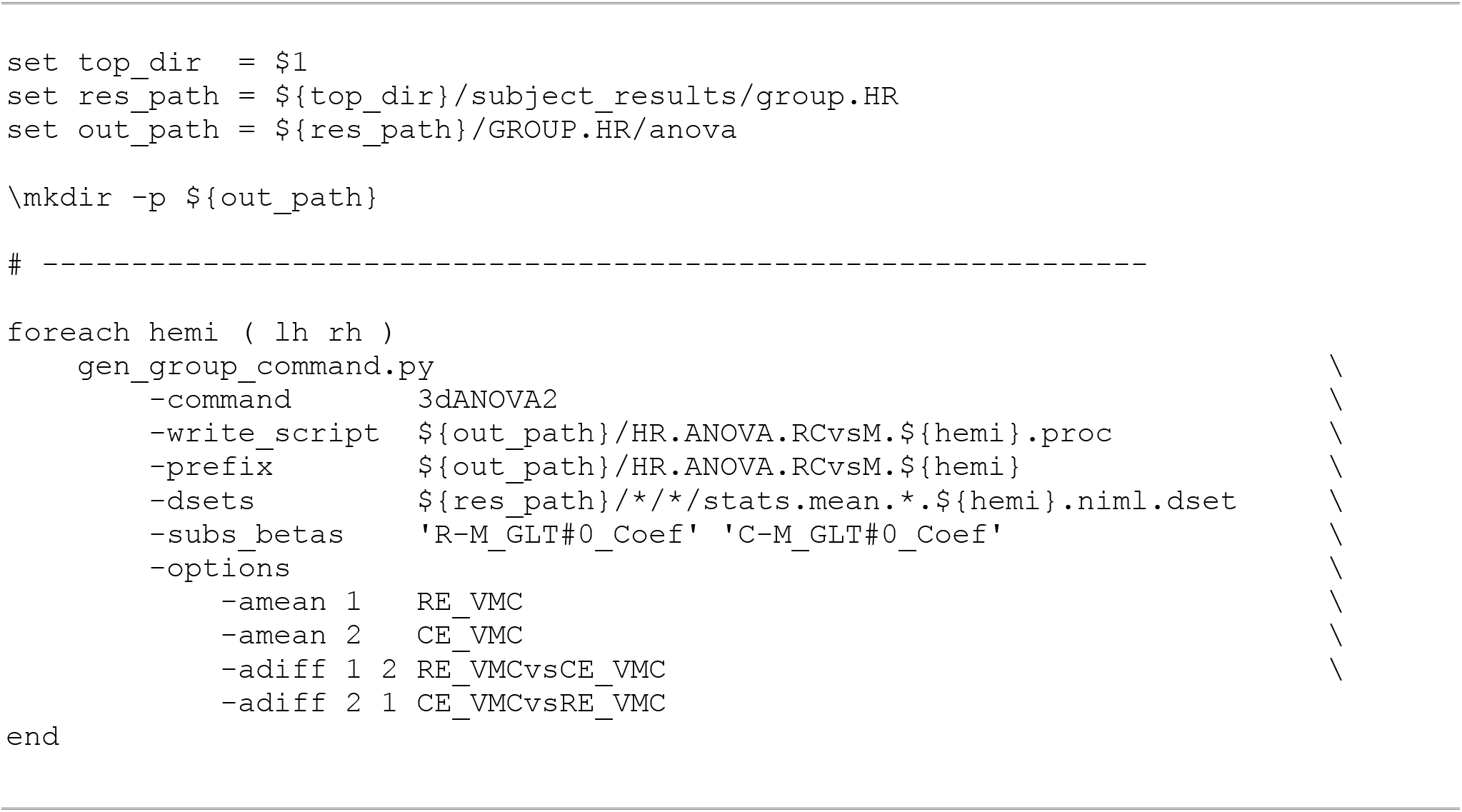
The command to generate ANOVAs for group analysis to observe the whole-brain activity pattern of CE and RE tasks in the present study. The following abbreviations are used: lh = left hemisphere; rh = right hemisphere; CE = curve estimation conditions; RE = roughness estimation conditions; VMC = visual motion control condition.

**Table S5.**
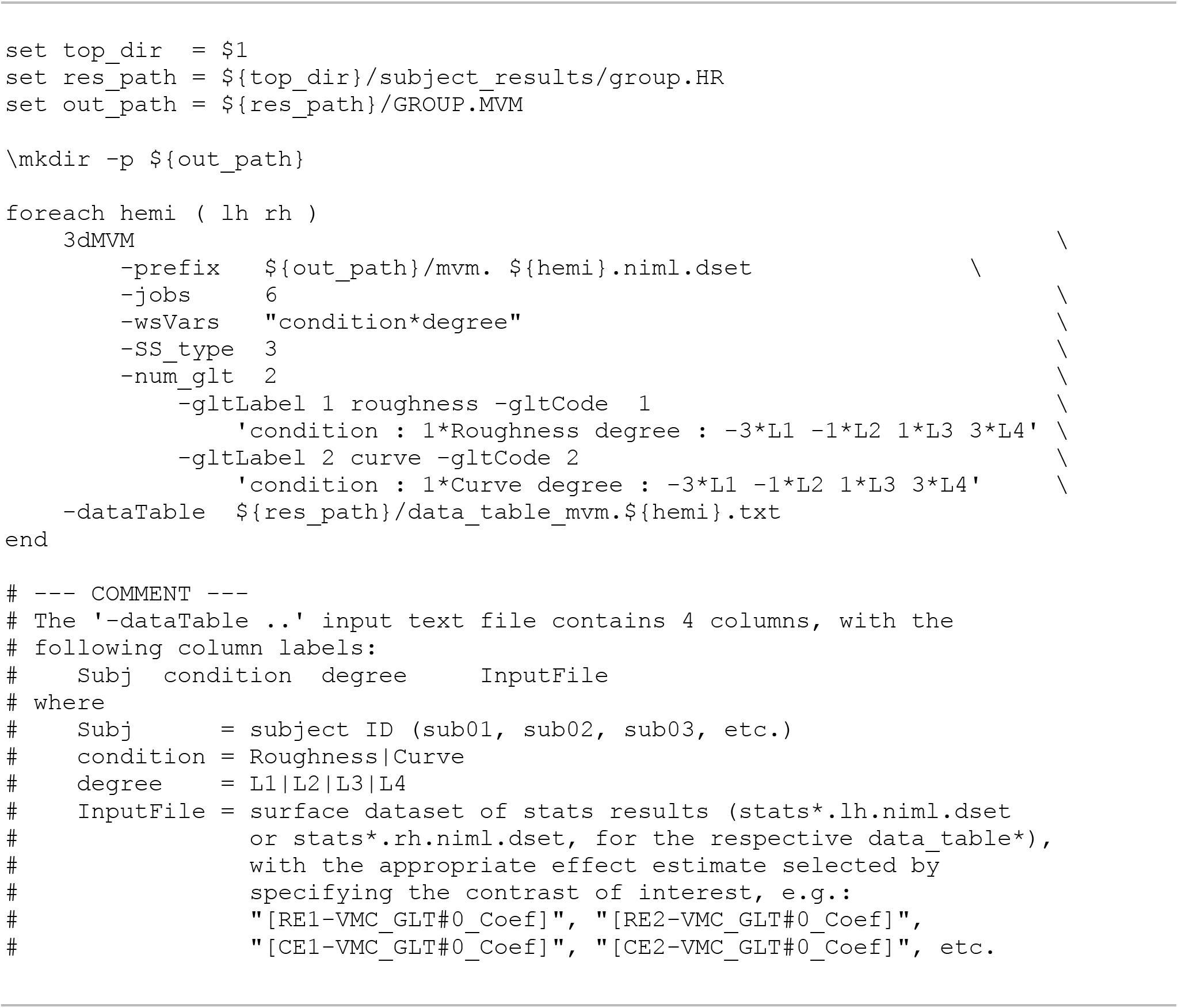
The command for MVM analysis with 3dMVM for group analysis to observe the brain regions parametrically modulated by CE and RE tasks in the present study. The following abbreviations are used: lh = left hemisphere; rh = right hemisphere; CE = curve estimation conditions; RE = roughness estimation conditions; VMC = visual motion control condition; L1 = degree level one (and similar for L2, etc.).

